# First Report of Soft Rot Caused by *Pectobacterium brasiliense* on Summer Squash in Mississippi, USA

**DOI:** 10.64898/2026.01.26.701841

**Authors:** Lewis Brooks, Prachi Bista, Emmanuel Clark, Frank Mrema, Bed Prakash Bhatta

**Affiliations:** Department of Agriculture, Alcorn State University, 1000 ASU Drive, Lorman, Mississippi 39096, USA; College of Science, Engineering & Technology, Jackson State University, Jackson, Mississippi 39217, USA

**Keywords:** *Cucurbita pepo*, zucchini, bacterial plant disease

## Abstract

Summer squash (*Cucurbita pepo*) is a popular vegetable in Mississippi. These are harvested during the tender and immature stages. This vegetable is known to be a good source of vitamins A and C, as well as potassium, and iron. Small farms typically sell summer squash directly to consumers through local farmers’ markets. In this study, we isolated and identified a soft rot causing bacteria, *Pectobacterium brasiliense* strain 25ASUB12 (GenBank: PX884501), from symptomatic fruit of field-grown summer squash in Mississippi. We deployed both phenotypic and molecular techniques to identify this important pathogen which has a wide host range, including cucumbers, potatoes, and tomatoes.

## Introduction

Summer squash (*Cucurbita pepo* L.), an important vegetable crop belonging to the gourd family Cucurbitaceae, is native to the Americas (Cutler & Whitaker, 1961) with evidence of domestication in the lower Mississippi River (Hart, 2008). *Pectobacterium brasiliense*, a highly virulent bacteria with pectinolytic abilities, was first reported to cause blackleg of potato in Brazil (Duarte et al., 2004). This study was conducted to isolate/identify a soft rot causing bacterial pathogen from symptomatic squash fruit, confirm its pathogenicity, and reisolate/identify to fulfill the Kochs’ postulates.

## Materials & Methods

### Isolation of Bacteria from Symptomatic Fruit Tissue

In July 2025, soft rot symptoms were observed on fruit of summer squash cv. Black Beauty grown at the Alcorn Experiment Station, Mississippi (31° 52’ 23.412”N 91° 7’ 46.739”W). The incidence of the disease was c. 20% (n =5). The outside of the fruit had mushy, water-soaked spots with visible oozing of the fruit tissue. The interior of the fruit showed wet, rotten, slimy brown tissue (**Figure 1**).

**Figure 1.**
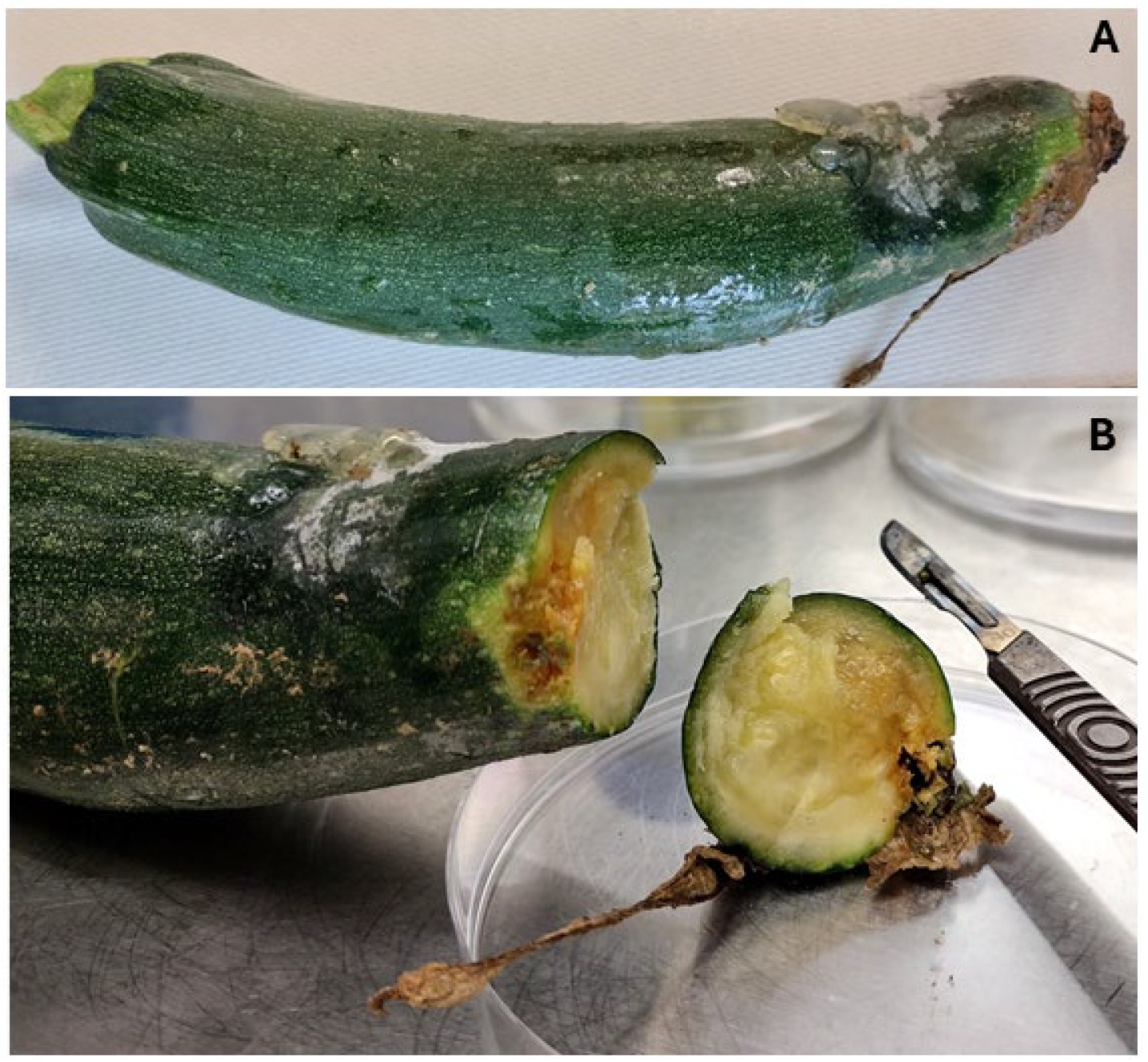
Symptoms of bacterial soft rot caused by *Pectobacterium brasiliense* on summer squash, (A) symptomatic whole fruit, and (B) the soft rot area on the cut fruit depicted by wet, slimy brown tissue.

Small section (5 mm x 5mm) of internal fruit tissue was excised from the margin of symptomatic areas and macerated in a microcentrifuge tube containing 100 µL of sterile phosphate-buffered saline (PBS, 1X). The bacterial suspension was streaked on nutrient agar (NA) medium and incubated at 25°C in the dark. A single colony was streaked on NA to obtain a pure culture of the bacterial strain designated as 25ASUB12 (**Figure 2**).

**Figure 2.**
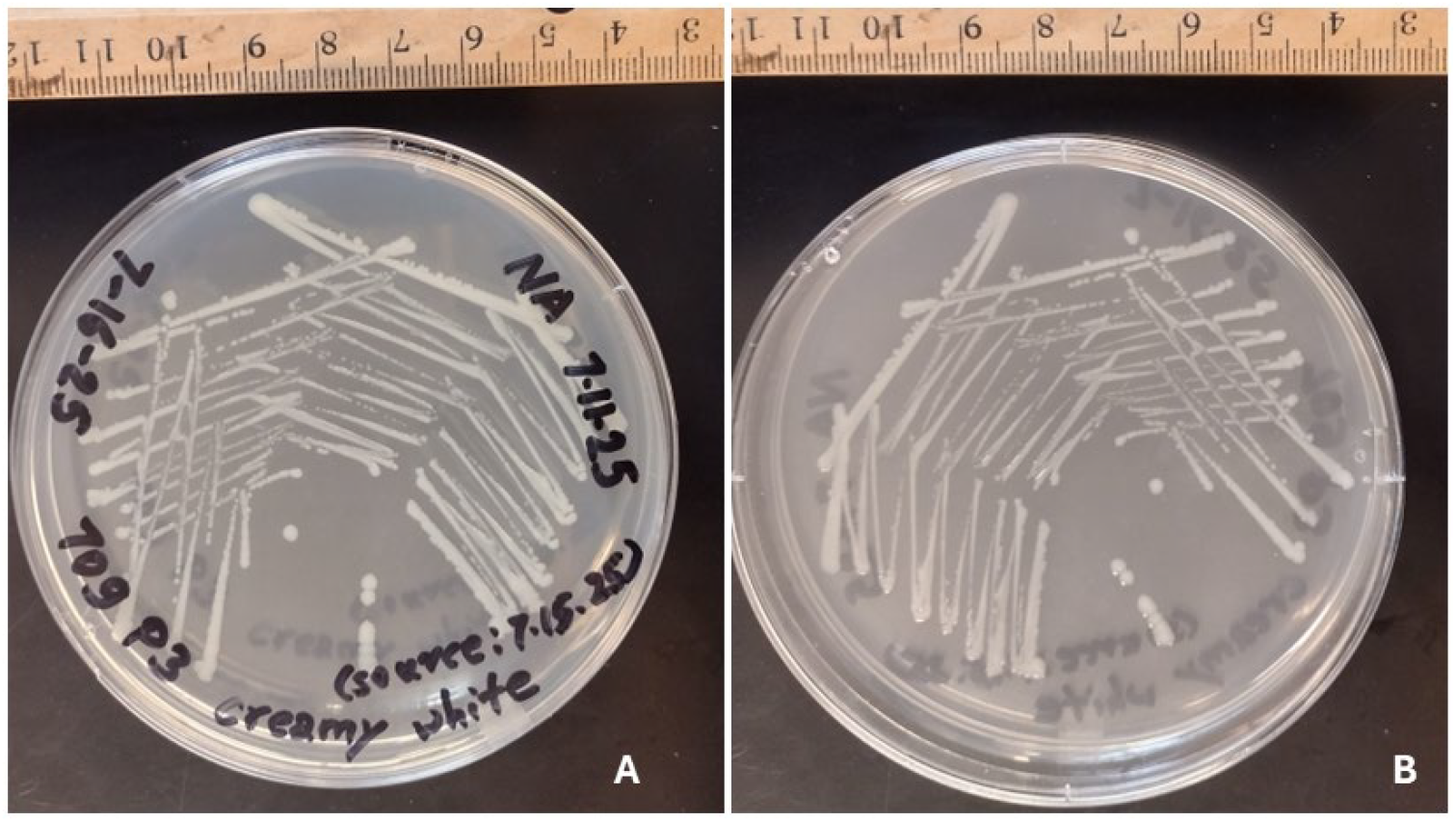
Pure culture of soft rot pathogen, *Pectobacterium brasiliense* strain 25ASUB12, isolated from summer squash fruit. (A) shows the bottom view of the bacterial colonies on nutrient agar and (B) shows the top view of the bacterial colonies on nutrient agar one day after streaking from the initial single colony.

### Molecular characterization of purified bacterial strain

Genomic DNA was extracted from two-day old bacterial culture grown on NA media using Quick-DNA Fungal/Bacterial Miniprep Kit (Zymo Research Corp., California, USA). The primers, 27F/1492R, were utilized to amplify the 16S ribosomal RNA (rRNA) gene. The PCR product was visualized via gel electrophoresis, purified using E.Z.N.A. Cycle Pure Kit (Georgia, USA), and sent for sequencing to Plasmidsaurus Inc., Kentucky, USA.

### Pathogenicity assay of *Pectobacterium brasiliense* strain 25ASUB12

Pathogenicity assay was conducted on summer squash fruits (cv. Black Beauty). Surface disinfection was done by spraying 70% ethanol on the summer squash fruits and wiping with sterile paper towel. Bacterial inoculum of *Pectobacterium brasiliense* strain 25ASUB12 was prepared from two-day old culture grown on NA (OD_600_ = 0.2). A 500 µL bacterial suspension was injected on the summer squash fruits using a sterile syringe and needle. The control fruits were inoculated with 500 µL of sterile PBS. These fruits were kept in a tray containing sterile paper towels moistened with sterile water and covered loosely with a lid for four days in the dark at 25°C.

## Results

### Molecular identity of Pectobacterium brasiliense strain 25ASUB12

The GenBank accession no. for 16S rRNA gene of *Pectobacterium brasiliense* strain 25ASUB12 is PX884501. This strain had >99% nucleotide identity (BLASTn; ‘core_nt’ database) to over thirty strains of *P. brasiliense*.

### Results from the pathogenicity assay

The treated fruits exhibited symptoms like those collected from the field while the control did not (**Figure 3**). The re-isolated daughter strains matched the colony morphology of the mother strain. Re-isolation from the inoculated fruits and sequencing resulted in six daughter strains of *P. brasiliense* strain 25ASUB12 (GenBank Accession Nos.: PX884502, PX884503, PX884504, PX884505, PX884506, and PX884507).

**Figure 3.**
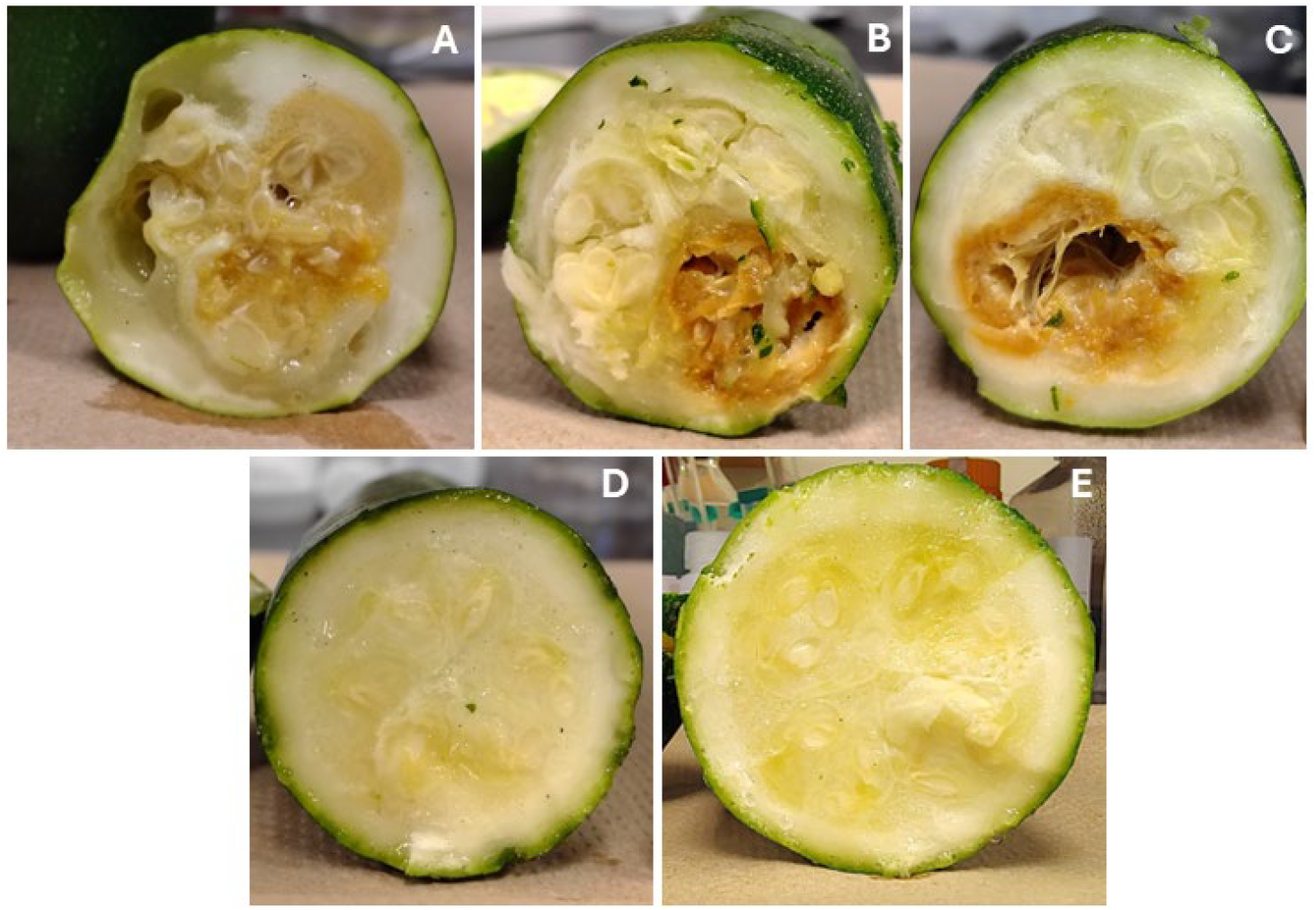
Pathogenicity of *Pectobacterium brasiliense* strain 25ASUB12 on summer squash cv. Black Beauty fruits; A, B, and C show the symptoms on summer squash fruits inoculated with bacterial suspension 4-days after inoculation; D, E show the summer squash fruits inoculated with control (sterile, 1X phosphate-buffered saline).

## Conclusion

To the best of our knowledge, this is the first documented case of *Pectobacterium brasiliense* causing soft rot of summer squash in Mississippi, USA. *P. brasiliense* has previously been reported as a pathogen of kale (Boluk et al., 2020), mizuna (Klair et al., 2021), pak choi (Klair et al., 2021) and potato (Zhang et al., 2023) in the USA. This report adds summer squash to the diversity of crops affected by *P. brasiliense*. Further investigation on prevalence of this soft rot disease in the region and development of disease management practices will be warranted.

## Acknowledgements

This research was supported by the intramural research program of the U.S. Department of Agriculture, National Institute of Food and Agriculture, Evans-Allen.

## Author Contributions

***Conceptualization***: B.P.B.; ***Methodology***: L.B., P.B., F.M., and B.P.B.; ***Software***: L.B., P.B., and B.P.B.; ***Validation***: B.P.B.; ***Formal Analysis***: L.B., P.B., and B.P.B.; ***Investigation***: B.P.B.; ***Resources***: F.M. and B.P.B.; ***Data Curation***: B.P.B.; ***Writing—Original Draft Preparation***: L.B., P.B., E.C., F.M., and B.P.B.; ***Writing—Review and Editing***: F.M. and B.P.B.; ***Visualization***: B.P.B.; ***Supervision***: B.P.B.; ***Project Administration***: B.P.B.; ***Funding Acquisition***: F.M. and B.P.B. All authors have read and agreed to the published version of the manuscript.

## Data Availability Statement

Data supporting the findings is available within the manuscript.

